# Spatially resolved transcriptomics reveals a unique disease signature and potential biomarkers for chronic traumatic encephalopathy

**DOI:** 10.1101/2025.04.23.650359

**Authors:** Catherine M Suter, Jennifer E Cropley, Andrew Affleck, Maggie Lee, Karina Hammond, Brian Gloss, Michael E Buckland

## Abstract

Chronic traumatic encephalopathy (CTE) is a poorly understood environmental tauopathy uniquely associated with repetitive head injury. It is clinically ambiguous, and at present can only be diagnosed post-mortem. There is a pressing need to understand CTE pathophysiology in order to identify targets for possible intervention and biomarkers for in-life diagnosis, however molecular characterisation of the disease is hampered by the stochastic distribution of CTE lesions. Here we have taken advantage of Visium spatial transcriptomics to map gene expression in discrete CTE lesions and matched normal tissue from the same individuals. In doing so we derived a common 21-gene “signature” of CTE lesions that highlights astrocytic activation, neuroinflammation, blood-brain barrier function, and extracellular matrix remodeling as key *in situ* features of CTE lesions. Almost all CTE signature genes were strongly expressed in astrocytes, and ontological and protein association analyses implicated extracellular matrix functions as drivers of the disease. These findings provide the first glimpse into the intricate molecular dynamics underlying CTE lesions *in situ*, and present 21 candidate molecules for the development of in-life CTE diagnostics.

## Introduction

Chronic traumatic encephalopathy (CTE) is a neurodegenerative disease uniquely associated with repeated exposure to traumatic brain injury (1). It has been linked to a syndrome that variously features cognitive and motor impairment, and neurobehavioural and mood dysregulation. However the syndrome is not specific to CTE, having significant overlap with other neurodegenerative conditions (2). CTE can currently only be diagnosed by postmortem histopathological examination of the brain.

CTE is defined by the accumulation of hyperphosphorylated tau protein (p-tau) in the form of neurofibrillary tangles inside perivascular neurons, and optionally astrocytes, located at the depths of the cortical sulci (3). CTE pathology exhibits a progressive spectrum extending from one or a few irregularly distributed p-tau foci in the cortex, through to brain-wide tauopathy (4). Since the precise location of CTE lesions is unpredictable, especially in early-stage disease, extensive brain sampling is often required for definitive diagnosis or exclusion.

There is a pressing need to better understand the molecular pathophysiology of CTE, so as to identify specific biomarkers, and develop methods for in-life diagnosis and treatment. But the unpredictable number, size and distribution of lesions – the same features that complicate CTE diagnosis – also confound its molecular interrogation.

Previous efforts to profile the molecular features of CTE have been hampered by the difficulty in knowing definitively whether a tissue sample contains a CTE lesion: bulk extraction of protein or RNA from a tissue sample precludes using that sample to confirm CTE pathology. Nevertheless, several studies have performed gene expression profiling on fresh-frozen samples of brains known to harbour CTE (5–10). While some gene expression changes have been identified, mainly in high stage disease, there is little concordance among studies, and in no study could gene expression changes be ascribed to the CTE lesion itself.

Spatial transcriptomics provides a transformative means of understanding the molecular underpinnings of CTE. It allows precise mapping of gene expression across a tissue section, and because it can be performed on material that has been fixed for histopathology, facilitates the transcriptional profiling of *bona fide* lesions that have been confirmed by p-tau immunohistochemistry. Thus, spatial transcriptomics provides a way to overcome the challenge that random distribution of CTE lesions presents to gene expression analysis.

We made use of the Visium spatial transcriptomics platform to map gene expression within CTE lesions and in nearby regions containing little to no p-tau. By doing this we uncovered a gene expression signature of CTE lesions that provides insight into disease pathophysiology as well as identifying potential diagnostic biomarkers. Collectively, the gene expression changes we observed point to astrocyte activation as a key feature of CTE pathology.

## Results

### Spatial transcriptomics of CTE lesions

We employed the 10x Genomics Visium platform to examine gene expression within and around CTE lesions from human frontal cortex. Visium combines transcriptome-wide sequencing with histological features to enable evaluation of gene expression within the context of tissue architecture (**Figure 1a**). Tissue sections are placed onto a Visium slide composed of four 6.5 mm x 6.5 mm grids of ∼5,000 ‘spots’; each spot is 55 µm in diameter and contains barcoded oligo-dT primers that capture mRNA from the tissue section. Reverse transcription and PCR is used to generate barcoded cDNA libraries that are sequenced and mapped back to their spot of origin. For CTE, the ability of Visium to capture mRNA from FFPE material permits profiling of discrete CTE lesions that can be identified by IHC prior to molecular analysis. We selected CTE lesions based on size (< 3 mm), the integrity of derivative RNA, and the absence of any neurodegenerative pathology other than CTE (**Figure S1**). Tissue sections containing verified CTE lesions from four individuals (**Table S1**) were transferred to a Visium slide using CytAssist and processed using the standard 10x protocol. Three of the lesions provided sequencing data of sufficient quality to proceed with analysis. From these, we obtained a total of 642,194,961 reads from 13,648 Visium spots; average reads per spot (± SD) was 47,086 ± 12,073, and the median number of unique transcripts per spot was 6,175 ± 1,019.

**Fig. 1:**
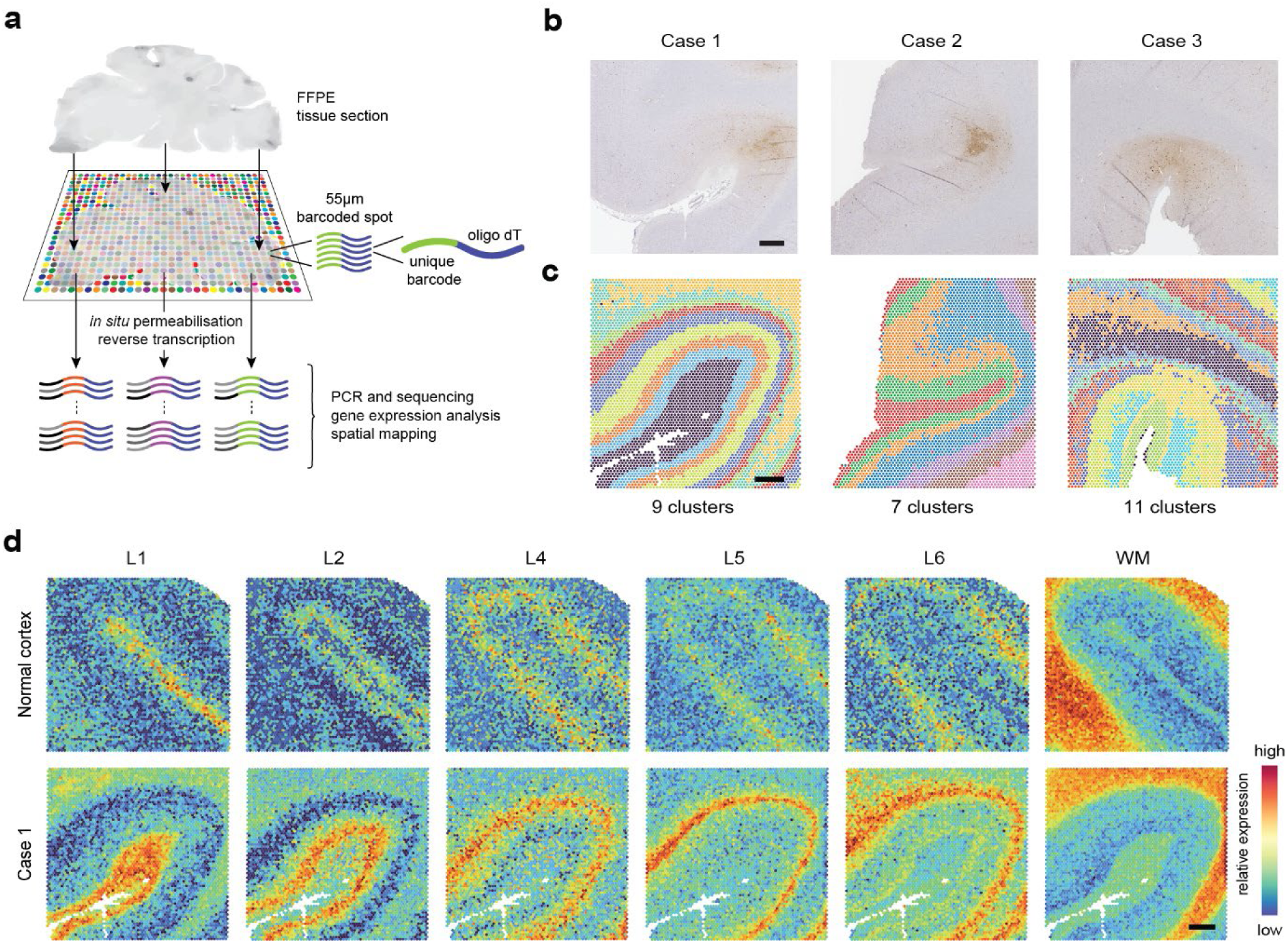
Visium reveals spatial patterns of gene expression that correspond to cortical layers. **a** Visium spatial transcriptomics involves placement of a formalin-fixed, paraffin-embedded (FFPE) tissue section on a Visium slide dotted with barcoded oligo dT primers, which bind to the RNA in the tissue. After *in situ* reverse transcription, the resulting barcoded cDNA is amplified by PCR and sequenced; sequence data is mapped back to the tissue section bioinformatically. **b** CTE cases 1-3. These tissue sections are serial to those used for Visium; IHC for p-tau (brown) shows CTE lesions at the depths of cortical sulci. Scale bar, 1 mm. **c** Unsupervised clustering of mapped Visium transcriptomes shows gene expression clusters that resemble the spatial patten of cortical layers. Scale bar, 1 mm. **d** Annotation of cortical layers of a reference normal cortex dataset (top row) and a representative CTE case (bottom row). L, cortical layer; WM, white matter.

To match transcriptional profiles to CTE lesions, we aligned the coordinates of each sample’s Visium grid to a serial section stained with AT8 (see Methods; **Figure 1b**). Unsupervised clustering of transcriptomes from all spots on the grid identified between 7 and 11 spatial gene expression clusters for each case (**Figure 1c**), but no cluster corresponded to the p-tau immunoreactive regions indicative of the CTE lesion. We did however notice that the expression clustering resembled cortical layer distribution, and thus applied previously described layer-specific gene modules (11) to annotate the cortical layers in our samples, as well as in a reference Visium dataset from normal human frontal cortex (12); **Figure 1d**. All the cortical layers were represented in each sample, and in a pattern that largely matched the unsupervised expression clustering. The only point of difference was an expansion of the first cortical layer in CTE cases relative to normal cortex (**Figure 1c; Figure S2**). We considered that the strong cortical layer gene expression signatures might be masking a CTE lesion-specific signature.

### Highly localised gene expression changes in CTE lesions

We proceeded to address the potential masking of CTE gene expression by subsetting the data spatially: for each case, we selected a subset of spots located within regions of dense p-tau immunoreactivity (i.e. within the CTE lesion) and compared the transcriptomes with those from adjacent areas with matched cortical layer representation but little to no p-tau (**Figure 2a**).

**Fig. 2.**
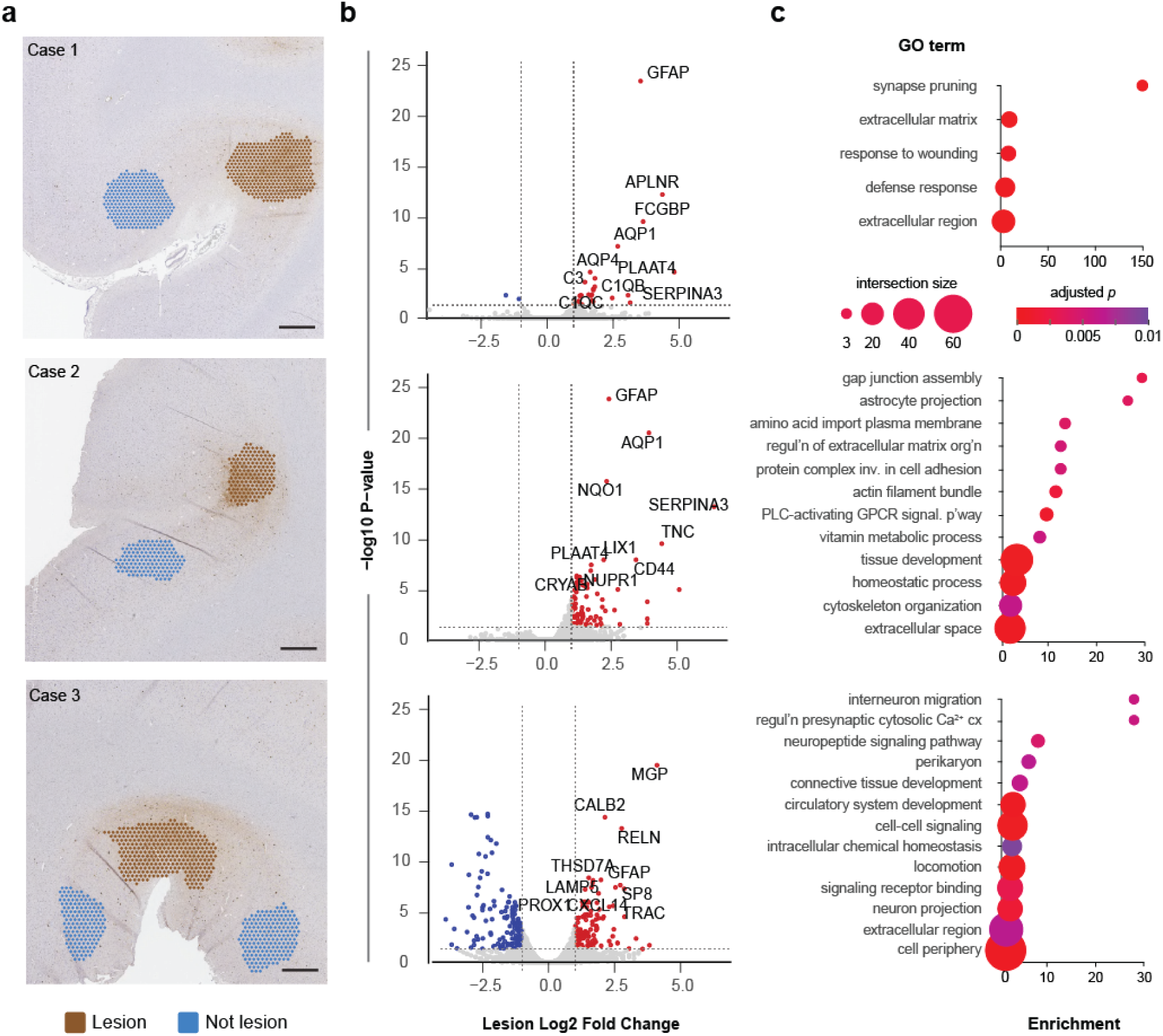
Genes upregulated in CTE lesions are involved in ECM functions. **a** For interrogation of CTE lesion-specific gene expression, serial tissue sections immunostained for p-tau (as shown in Figure 1) were used to map Visium spots corresponding to CTE lesions (Lesion, brown) and to nearby areas with no p-tau but matched cortical layer gene expression (Not lesion, blue). Scale bar, 1 mm. **b** Scatter plots showing gene expression differences between Lesion and Not lesion. Dotted lines mark a fold-change threshold of > 2, and FDR threshold of < 0.05. Upregulated genes are shown in red, and downregulated genes in blue. **c** Top gene ontology (GO) terms for genes upregulated in CTE lesions. Only terms with Benjamini-Hochberg corrected *p*-value < 0.01 and more than two upregulated genes represented (i.e. intersection size ≥ 3) are shown.

Differential gene expression analysis showed 32, 102, and 130 upregulated transcripts from cases 1, 2, and 3, respectively (using stringent criteria of FC > 2, FDR < 0.05; **Figure 2b**). In all three samples, these upregulated transcripts shared ontologies related to extracellular matrix (ECM) functions (**Figure 2c**). The full list of upregulated genes and enriched ontologies for each sample are detailed in **Table S2**.

Surprisingly, lesions in case 1 and 2 exhibited very few downregulated transcripts in the CTE lesion compared to adjacent tissue. Only case 3 exhibited a substantial number of downregulated genes (**Figure 2**). These transcripts did not cluster into any functional categories, so their significance is unclear. It is worth noting that the CTE lesion in case 3 is more diffuse than in the other cases (**Figure 1 and S1**), and so it is possible that the larger number of dysregulated transcripts represent a more advanced lesion with gene expression changes secondary to disease progression.

### A common transcriptomic signature implicates astrocytes in CTE

Because we were seeking a positive signature of CTE to identify potential biomarkers, we directed our attention to genes upregulated in CTE lesions relative to adjacent cortex. We focused on common elements across the three cases, where at least two showed statistically significant upregulation in the lesion. We found 21 genes that fit these criteria (**Figure 3a**). While most of these (17/21) were statistically upregulated in only two of three cases, the third case invariably exhibited the same directionality of change, and a similar magnitude of change (**Figure 3b, Table S2**). These 21 genes provide a potential transcriptomic signature of CTE.

**Fig. 3.**
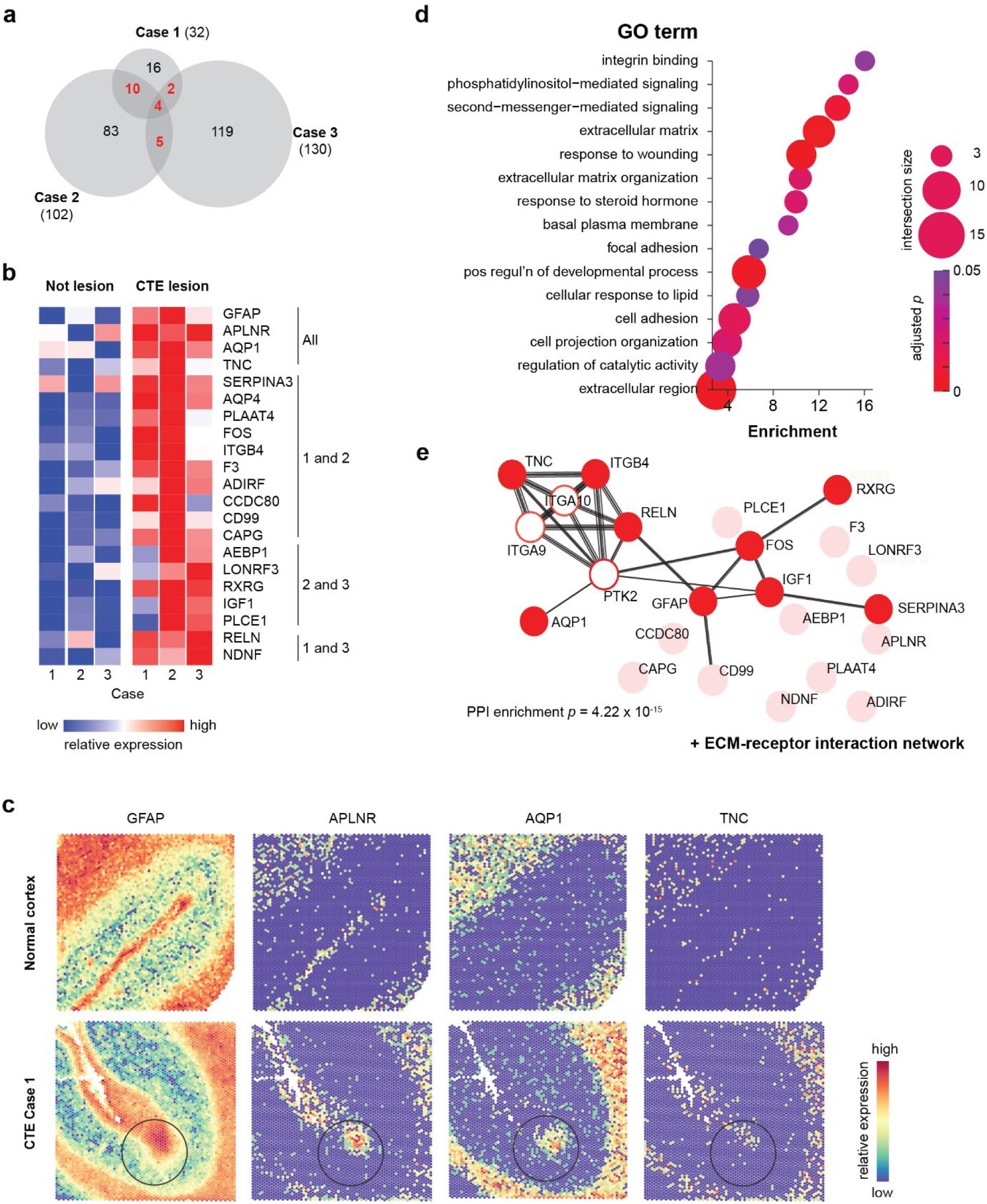
Genes commonly upregulated in CTE lesions reveal a CTE expression signature. **a** Venn diagram showing overlap of genes upregulated in lesions across the three CTE cases. **b** Gene expression heatmap of the 21 genes upregulated in lesions of at least two of the three CTE cases. **c** Visium spatial maps of expression of the four genes upregulated in lesions of all three CTE cases, in a normal cortex reference dataset (top row) and a representative CTE case (bottom row). The black circle shows the location of the CTE lesion. **d** Top GO terms for the 21 signature genes. Only terms with Benjamini-Hochberg corrected *p*-value < 0.05 and more than two upregulated genes represented (i.e. intersection size ≥ 3) are shown. **e** The top network from an unbiased protein-protein interaction (PPI) analysis, with signature genes in red. Signature genes not in the network are shown in pink; interactor proteins not in the signature gene list are shown in white.

Of the 21 genes, there were four that were significantly upregulated in all three CTE lesions: *GFAP*, *APLNR*, *AQP1* and *TNC* (**Figure 3c**), implying that astrocytic activation, neuroinflammation, blood-brain barrier compromise, and ECM remodeling are universal features of CTE pathophysiology. Taken together, the 21 CTE signature genes were enriched for multiple ontologies (**Figure 3d; Table S2)**, and again, functions relating to the ECM were highly significant. Furthermore, unbiased protein association analysis using the STRING database (13) found that nine of the 21 CTE genes formed a network of known ECM-receptor interactions (**Figure 3e**).

The resolution of Visium (55 µm spots, with 45 µm gap between spots) does not permit gene expression changes to be assigned to single cells. To understand which cell types express the CTE signature genes, we interrogated publicly available single nucleus RNAseq data (14). We found that the majority of the 21 genes were expressed predominantly in astrocytes, with other non-neuronal cells, including vascular and leptomeningeal cells and endothelial cells, also prominent; notably, few of the genes were expressed predominantly in neurons (**Figure S3, Table 1**). We also queried the metadata-derived ‘AD gene expression portrait’ (ADp; (15)) to find that all but two of the 21 CTE signature genes are also upregulated in AD (**Table 1**), although not necessarily to the same extent. Taken together, these data suggest that CTE shares molecular features with Alzheimer’s Disease, and that CTE appears largely driven by non-neuronal processes.

**Table 1:** Cell type expression profiles of the 21 gene CTE signature.

| **Gene** | **Main cell type expression<sup>a</sup>** | **ADp expression (rank)<sup>b</sup>** |
| --- | --- | --- |
| ADIRF | A, EC, VLMC | ↑ (n/a) |
| AEBP1 | A, VLMC | ↑ (6) |
| APLNR | A, VLMC | ↑ (549) |
| AQP1 | A, VLMC | ↑ (777) |
| AQP4 | A | ↑ (206) |
| CAPG | A, M-PVM, | ↑ (492) |
| CCDC80 | A, L5/6 NP, VLMC | ↑ (n/a) |
| CD99 | A, EC, M-PVM, VLMC | ↑ (261) |
| F3 | A, VLMC | ↑ (n/a) |
| FOS | A, EC, M-PVM, O, OPC, VLMC | ↑ (n/a) |
| GFAP | A | ↑ (2) |
| IGF1 | M-PVM, N | Decreased (n/a) |
| ITGB4 | A, O, OPC | ↑ (470) |
| LONRF3 | ubiquitous | ↑ (n/a) |
| NDNF | N | ↑ (n/a) |
| PLAAT4 | A, EC, M, O | ↑ (n/a) |
| PLCE1 | A, N, OPC, VLMC | ↑ (28) |
| RELN | N | No change (n/a) |
| RXRG | A, N | ↑ (n/a) |
| SERPINA3 | A, VLMC | ↑ (176) |
| TNC | A | ↑ (n/a) |

<sup>a</sup> Data from (14); <sup>b</sup> Direction of expression in Alzheimer disease gene portrait (ADp) and ranking of top 1000 genes upregulated in AD from (15); A, astrocyte; EC, endothelial cell; M-PVM, microglia-perivascular macrophage; N, neuron; NP, pyramidal neuron; O, oligodendrocyte; OPC, oligodendrocyte precursor cell; VLMC, vascular and leptomeningeal cell.

### Insights from immunohistochemical validation

Of the four genes we identified as universally upregulated in CTE lesions, two – *AQP1* and *GFAP* – have been previously linked to CTE (10). We thus examined the relationship between the spatial transcriptional changes of these genes and corresponding spatial protein expression using IHC. We included the three CTE cases used for Visium plus an additional five CTE cases (also from individuals with no other neurodegenerative pathology), as well as three control cases with no neurodegenerative pathology. We found that seven of the eight CTE cases had increased expression of both proteins relative to controls (**Table S1, Figures 4 and 5**), confirming that the elevated transcript levels observed in Visium conferred an increase in protein.

**Fig. 4.**
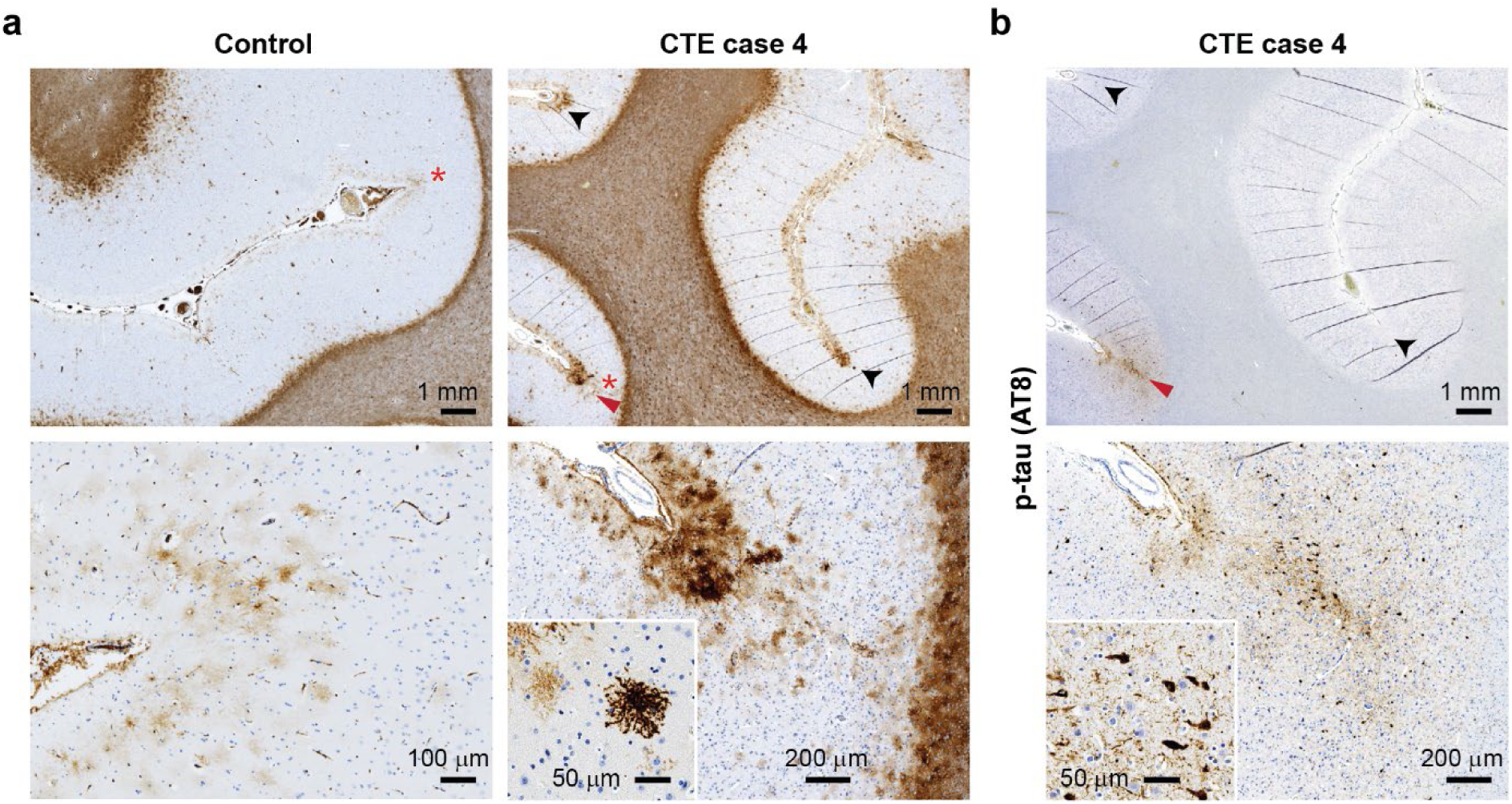
Representative immunohistochemistry of aquaporin 1 (AQP1) expression in CTE. **a** IHC for AQP1 in a control case (left panels) and CTE case 4 (right panels), at low (top) and high (bottom) magnification. Asterisks denote zoomed regions. The CTE lesion is marked with a red arrowhead, as defined in (**b**) a serial section of case 4 stained for p-tau. Note AQP1 accumulation at p-tau negative sulcal depths as well as in the CTE lesion (black arrowheads). Insets show individual cell morphology. Scale bars as shown.

**Fig. 5.**
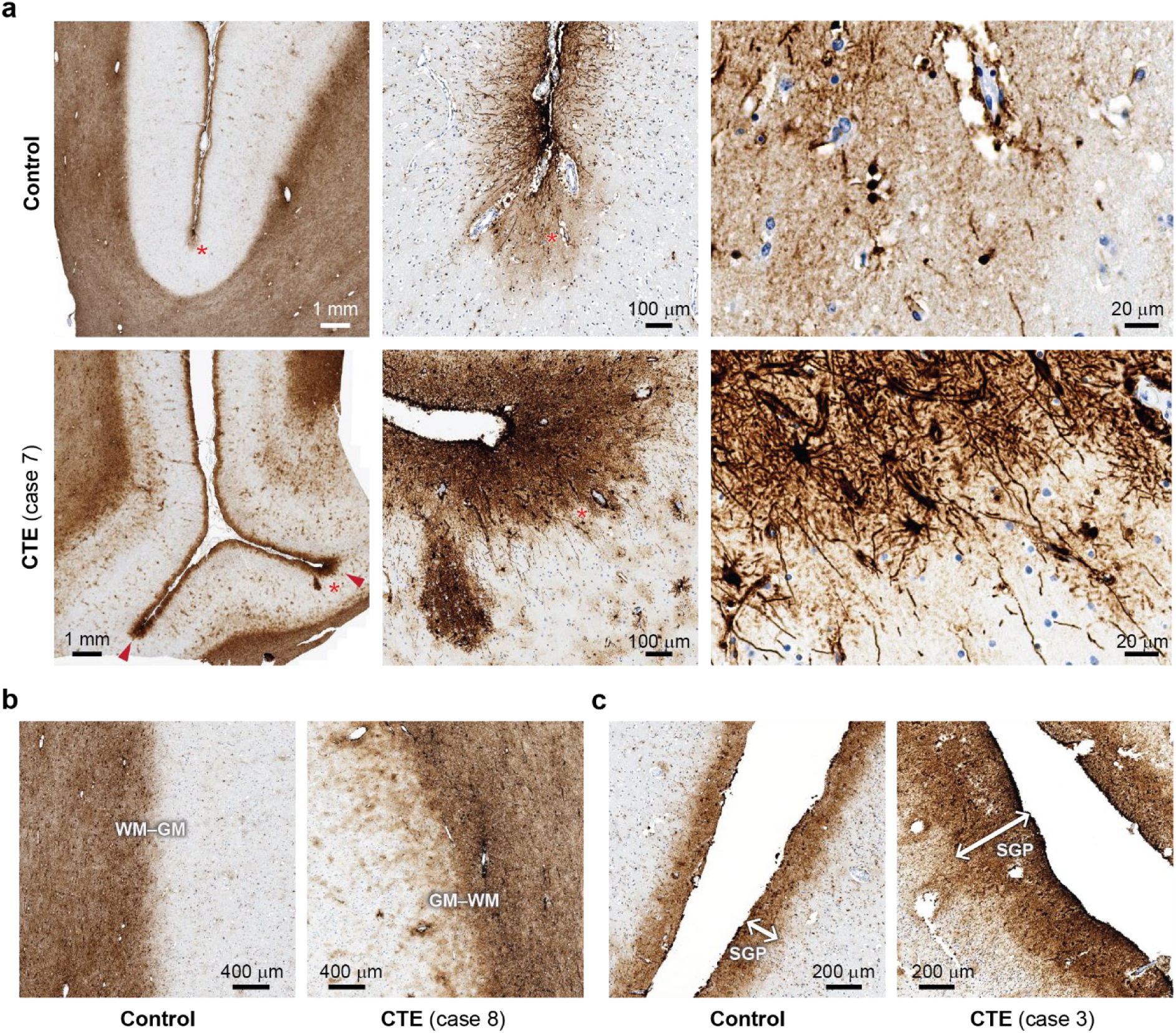
Representative immunohistochemistry of glial fibrillary acidic protein (GFAP) expression in CTE. **a** GFAP expression in a control case (top panels) and CTE case 7 (bottom panels), at low (left), medium (middle) and high (right) magnification showing a depth of sulcus (red asterisks mark magnified regions); locations of CTE lesions are marked with red arrowheads. Note higher expression of GFAP in grey matter in CTE, along with an increase in size and number of astrocytes, indicative of reactive astrogliosis. **b** GFAP expression at the grey-white matter junction, in a control case (left) and CTE case 8 (right). Note GFAP expression extending into grey matter and blurring the grey-white matter junction. **c** GFAP expression in the subpial glial plate (SGP), in a control case (left) and CTE case 3 (right); note SGP thickening in the CTE case, reminiscent of interface astroglial scarring. Scale bars as shown.

Aquaporin 1 protein (AQP1) showed strong immunoreactivity in white matter in both control and CTE, but the CTE cases showed far denser staining at sulcal depths where CTE lesions were present (**Figure 4a, b**, red arrowheads). In three of the eight CTE cases, moreover, we also noted an accumulation of AQP1-positive cells at depths of sulci that did *not* harbour a CTE lesion, i.e., in areas with no p-tau (**Figure 4a**, black arrowheads). This raises the possibility that aberrant AQP1 expression precedes the accumulation of p-tau in CTE.

At high magnification, the AQP1-positive cells at the sulcal depths in CTE cases demonstrate variably dense immunoreactivity of cell bodies and ramifying cell processes typical of protoplasmic astrocytes (**Figure 4a**, inset). While there is overlap with the subpial p-tau distribution, AQP1 has a wider distribution and stains cells with distinctive protoplasmic astrocytic morphology, in contrast to thorn-shaped astrocytes of aging-related tau astrogliopathy (ARTAG) and neurofibrillary tangles (**Figure 4b**, inset).

Like AQP1, glial fibrillary acidic protein (GFAP) immunoreactivity is predominant in white matter and also in the glia limitans of the cortex in all samples (**Figure 5**). There was a noticeable difference in GFAP staining patterns in CTE cases compared to control. There was greater GFAP immunoreactivity at sulcal depths in CTE cases, in and around the CTE lesion (**Figure 5a**, left panels). This appears due to a greater number of GFAP-positive astrocytes compared to controls (middle panels); many of these astrocytes appear larger with processes increased in size and complexity, typical of reactive astrogliosis (right panels).

In four of the eight CTE cases with increased subpial astrogliosis, we also observed increased GFAP immunoreactivity outside of sulcal depths, around small intracortical blood vessels and at the grey-white junction (**Figure 5b**). These four cases also exhibited markedly thicker GFAP immunoreactivity in the subpial region (**Figure 5c)**. Similar observations have been made in association with interface astroglial scarring (IAS), a pattern of gliosis associated with blast-overpressure injury (16), and together indicate that features of IAS can accompany CTE in some individuals.

## Discussion

This study provides the first transcriptome-wide analysis of discrete CTE lesions, revealing a novel 21-gene signature associated with the disease. Our use of spatially resolved transcriptomics circumvented the challenges posed by the sporadic distribution of CTE lesions, which dilute any signal from bulk molecular interrogation techniques. The 21-gene signature not only represents a suite of candidate CTE biomarkers and potential therapeutic targets, but also provides fundamental insight into disease pathophysiology.

The 21-gene CTE signature shows a strong association with ECM remodeling and astrocytic processes. ECM-related ontologies were overrepresented in the signature, and also in upregulated genes from individual lesions. Functional protein-protein association analysis revealed one highly significant network – of ECM-receptor interactions – which contained nine of the 21 genes. In terms of astrocytic involvement, the overwhelming majority (18/21) of signature genes are expressed in astrocytes, many predominantly so. These findings extend prior observations of astrocytic activation and degeneration in CTE (17), as well as upregulation of ECM-related genes in bulk preparations of brains with high-stage CTE (10). Together these suggest that interplay between astrocytes and the ECM is pivotal in the development and progression of CTE.

Two of the four genes upregulated in all three CTE lesions – *GFAP* and *AQP1* – have previously been implicated in CTE (10). Immunohistochemical examination of GFAP and AQP1 protein expression revealed a close association with CTE lesions. Both proteins exhibited increased expression in p-tau positive sulcal depths in seven of the eight cases used for validation, confirming our transcriptomic findings at the protein level, and in a larger cohort.

AQP1 was prominently expressed in astrocytic processes where CTE lesions were present, even though the CTE lesions in this study displayed primarily neuronal p-tau. Notably, in some cases we observed AQP1-positive astrocytes in sulcal depths without *any* p-tau. This suggests that aberrant AQP1 expression may precede tau pathology in CTE, providing a promising early biomarker for diagnosis. Aberrant AQP1 expression, particularly in the absence of p-tau pathology, supports the hypothesis that astrocytic dysregulation may be an early driver of CTE lesion formation. The 21-gene signature also included a second brain aquaporin, *AQP4*, well known to be perturbed in AD (18). Taken together with the prominence of vasculature-related gene expression changes, this suggests that maintenance of blood-brain-barrier (BBB) integrity is impaired early in the development of CTE lesions. BBB compromise would allow entry of inflammatory mediators from the periphery, contributing to the progression of disease. Such compromise might however present an advantage for CTE detection: any astrocyte-derived biomarkers released into the periphery through the compromised BBB could potentiate non-invasive, in-life diagnosis of CTE.

We found GFAP highly expressed in and around CTE lesions, along with astrocytic hypertrophy and process elaboration, together indicating reactive astrogliosis. We also observed thickened GFAP-positive astrocytic barriers at the grey-white matter junction and subpial regions in four of the eight CTE cases used for validation. These are features similar to those described in IAS, previously reported in blast-induced traumatic brain injury (16), and in others with a history of repetitive sport-related head injuries (both with and without CTE) (19). Our data indicate that IAS may accompany CTE, but that is it not a universal feature of the disease.

Perhaps unsurprisingly, many genes in the CTE signature overlapped with genes dysregulated in the most common tauopathy, AD. Of the 21 signature genes, 19 have been identified as consistently upregulated in AD, with two, *AEBP1* and *GFAP*, highly upregulated (15). Both of these have been proposed as candidate blood-based AD biomarkers (20, 21) but our data suggests that they would not be specific for AD. By contrast, the 21-gene CTE signature provides candidates for CTE biomarkers that may distinguish it from other tauopathies such as AD. For example, *IGF1*, a key regulator of neuronal survival (22), exhibits opposing expression patterns in CTE and AD: while upregulated in CTE lesions, *IGF1* is consistently downregulated in AD (15, 23). We also observed a robust increase in the expression of *RELN*, another factor integral to neuronal survival, which has recently been reported to be downregulated in AD (24). Moreover, its gene product Reelin has been identified as significantly upregulated in the serum of former athletes at high risk of CTE (25); taken together with our data, this highlights Reelin as an exciting potential CTE biomarker that could bring us closer to in-life diagnosis. From a pathophysiological viewpoint, the upregulation of *IGF1* and *RELN* suggests that CTE activates a transcriptional program of neuronal survival that may not occur in AD, at least when it has progressed to the cortex. It is also possible that p-tau structural polymorphism between CTE and AD (26) can trigger different survival pathways.

While this study has provided novel insights into the molecular landscape of CTE, there are limitations. The Visium technology available when this study was done did not allow for single-cell resolution, necessitating an indirect approach to cell type attribution (integration with public single-nucleus RNAseq data). It is likely that further insights would be gained with orthogonal data derived from the same samples. Multi-omics approaches on larger cohorts would be required to achieve this, and future studies should include other tauopathies to ascertain the disease specificity of the signature we have identified here. Additionally, since Visium technology is designed for intra-sample rather than inter-sample comparisons, we did not compare CTE tissue with equivalent samples that were CTE-negative, or RHI-naïve. While such comparisons may have revealed field effects, or pan-cortical changes due to CTE, our approach – comparing CTE lesions with matched pathologically normal tissue from the same sample – allowed us to identify changes specifically related to CTE pathology without confounding from inter-individual and compositional factors.

## Conclusion

This study significantly advances the understanding of CTE by identifying a molecular signature that is disease-specific and pathologically grounded. The shared features between CTE and AD highlight the need for careful consideration of co-morbid tauopathies, but the unique non-neuronal drivers of CTE revealed here provide avenues for future research. The 21-gene CTE signature offers a promising foundation for in-life diagnosis of CTE. Such advancements are essential not only for accurately identifying affected individuals, but also for development and evaluation of therapeutic intervention. Without the ability to diagnose CTE during life, meaningful progress in treatment strategies will remain unattainable.

## Abbreviations

AD: Alzheimer’s Disease
ADp: Alzheimer’s Disease Portrait
APLNR: Apelin Receptor
AQP1: Aquaporin 1
ARTAG: Ageing-related tau astrogliopathy
BBB: Blood-brain barrier
cDNA: complementary DNA
CTE: Chronic Traumatic Encephalopathy
ECM: Extracellular Matrix
FC: Fold Change
FDR: False Discovery Rate
FFPE: Formalin-fixed Paraffin-embedded
GFAP: Glial Fibrillary Acidic Protein
GM-WM: Grey Matter – White Matter
GO: Gene Ontology
IAS: Interface Astroglial Scarring
IHC: Immunohistochemistry
PCR: Polymerase Chain Reaction
PPI: Protein-protein Interaction
p-tau: Hyperphosphorylated tau
RNAseq: RNA sequencing
SD: Standard Deviation
SGP: Subpial Glial Plate
TNC: Tenascin C

## Methods

### Tissue samples

This study was approved by the Sydney Local Health District Ethics Review Committee (Royal Prince Alfred Hospital (RPAH); X19-0010 and X23-0073) with informed consent from donors (prior to death) or their senior-next-of-kin. Clinical neuropathological evaluation was performed through the Australian Sports Brain Bank at the RPAH Department of Neuropathology (M.B.); formalin-fixed paraffin-embedded (FFPE) tissue was used in all analyses. Demographics of CTE cases (*n* = 8) and controls, who had a history of traumatic brain injury but no evidence of CTE (*n* = 3), are shown in Table S1.

### Immunohistochemistry

All immunohistochemistry was performed on a Leica Bond-MAX autostainer using the manufacturers’ recommendations with the following primary antibodies: AT8 (Thermo Fisher MN1020; 1:100), AQP1 (Thermo Fisher MA532593, 1:500), GFAP (Millipore MAB360, 1:400).

### Spatial transcriptomics

Three serial 5 µm sections were cut from FFPE tissue blocks from four CTE cases under RNAse-free conditions and placed onto glass slides. Slides were dried at 42 °C for three hours, and stored with desiccant until use. The first and third sections were subject to IHC with AT8 to verify the precise location of CTE lesions. The middle section was subject to Visium CytAssist spatial transcriptomics, performed in accordance with the manufacturer’s protocol (CG000542; 10x Genomics, Pleasanton, CA, USA) using the Human Probe Set version 2.0, at the South Australian Genomics Centre. Resultant cDNA libraries were sequenced using Illumina Nextseq High chemistry v2.

### Data preprocessing

Fastq files were mapped against the human genome hg38 (2020) and the Visium Human Transcriptome Probe Set version 2.0 using 10x Space Ranger (version 2.0.0) with manually aligned images from serial AT8 IHC. Reference data from normal human cortex were accessed from the 10x data repository (27). Subsequent analyses of mapped data were performed as below, either in 10x Loupe browser (28) (version 7.01), or with Seurat (29) in the R environment (version 4.3).

### Data analysis

Unsupervised spatial spot clustering was performed in Loupe using the graph-based clustering function. Cortical layer assignment of spots was performed according to Chen *et al.* (11). Differential expression analysis was performed between AT8-positive spots (Lesion) and AT8-negative spots (Non-lesion) within each CTE case in Loupe using default parameters. Only transcripts with greater than a 2-fold change and Benjamini-Hochberg corrected *p*-value < 0.05 were considered as differentially expressed in any further analyses. Gene ontology on differentially expressed genes was performed with gProfiler using genes within the Visium Probe set as background. Protein-protein interactions were assessed using STRING (13) with default parameters allowing only five or fewer non-query interactors.

## Declarations

### Ethics approval and consent to participate

This study was approved by the Sydney Local Health District Ethics Review Committee (Royal Prince Alfred Hospital (RPAH); X19-0010 and X23-0073). Written informed consent for brain donation was provided either by the donor prior to death, or the donor’s senior next-of-kin or executor.

### Consent for publication

Donors or next-of-kin provided consent for publication of their anonymised data.

### Availability of data and materials

The datasets used in this study are available from the corresponding author on request.

### Contributions

C.S. conceived and designed the study. A.A. was responsible for case curation. K.H, B.G. and C.S. were responsible for Visium experiments and analysis. M.L., A.A. and M.B. were responsible for immunochemistry and neuropathological characterisation. C.S. performed gene and protein association analysis. C.S., B.G., J.C. and M.B. interpreted data. C.S. and J.C. wrote the manuscript. All authors read and approved the final manuscript.

## Acknowledgements

All authors are grateful to the ASBB brain donors and their families who made this study possible. We also thank Brett Kennedy and James Fraser from 10x Genomics for assistance and advice, and the South Australian Genomics Centre (SAGC) for Visium CytAssist services; the SAGC is supported by the National Collaborative Research Infrastructure Strategy (NCRIS) via BioPlatforms Australia and by the SAGC partner institutes. Cell type attributions were based on data obtained from the AD Knowledge Portal (14); study data were generated from postmortem brain tissue obtained from the University of Washington BioRepository and Integrated Neuropathology (BRaIN) laboratory and Precision Neuropathology Core, which is supported by the NIH grants for the UW Alzheimer’s Disease Research Center (P50AG005136 and P30AG066509) and the Adult Changes in Thought Study (U01AG006781 and U19AG066567), and NIA grant U19AG060909.

