## Supplementary for "Spatially resolved transcriptomics reveals a unique disease signature and potential biomarkers for chronic traumatic encephalopathy": Figure S1.pdf

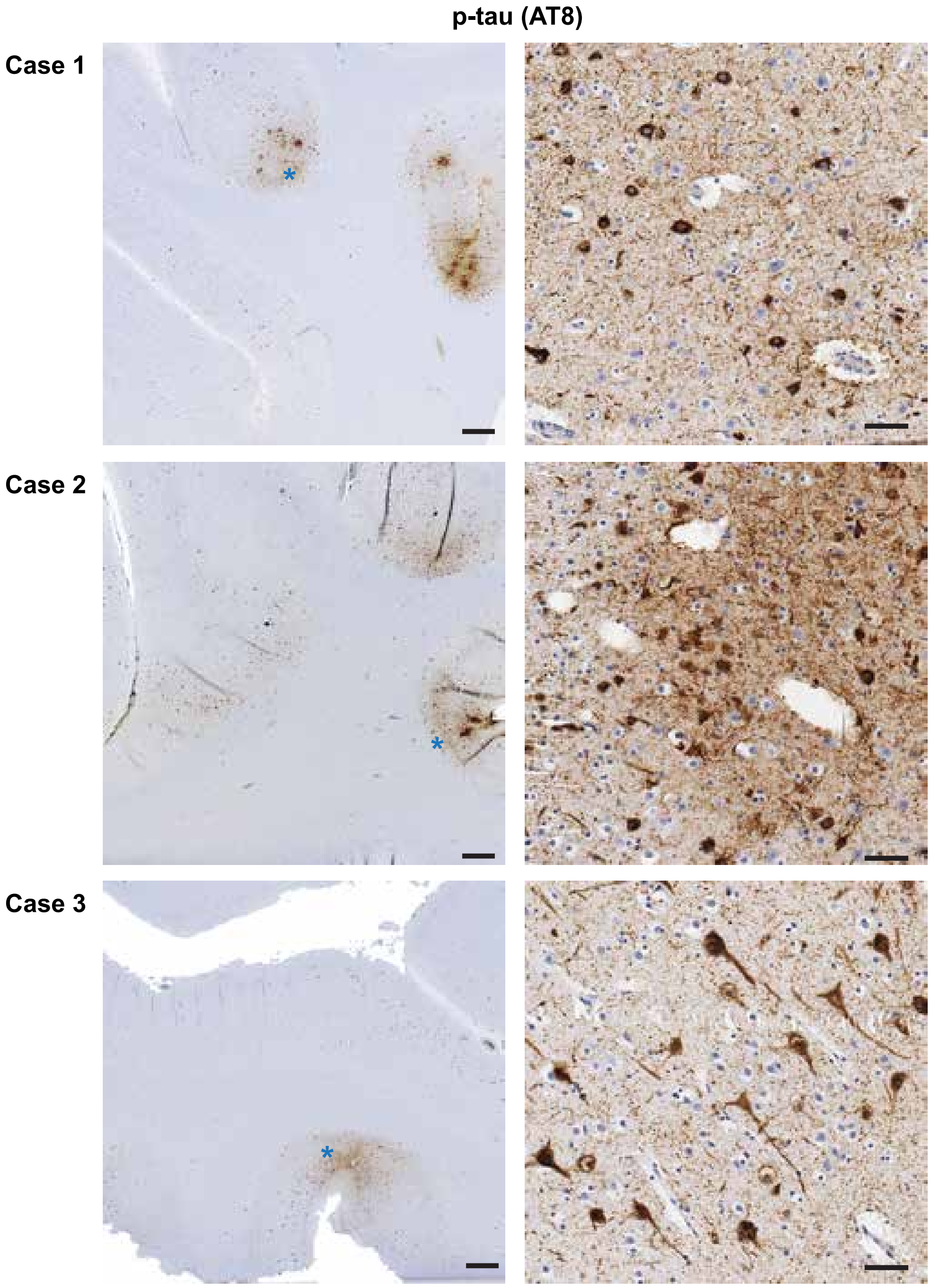


**Figure S1. AT8 immunohistochemistry of CTE cases used for Visium.**
The CTE lesions chosen for analysis are marked by the blue asterisk.
Scale bar: 1 mm for left panels, 50 μm for right panels.
