## Supplementary for "Spatially resolved transcriptomics reveals a unique disease signature and potential biomarkers for chronic traumatic encephalopathy": Figure S2.pdf

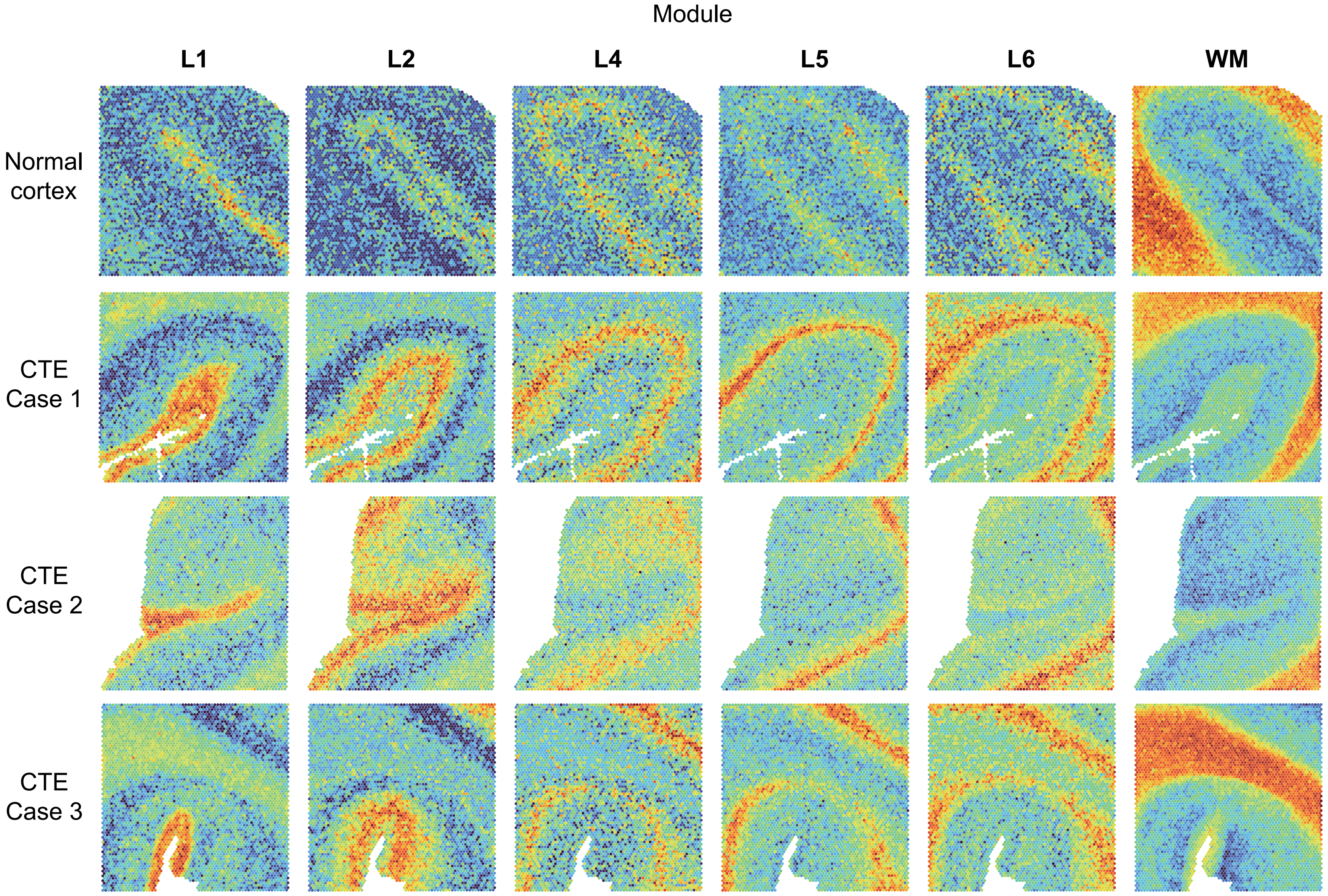


**Figure S2. Cortical layer distribution.** Shown are Visium transcriptomic heatmaps for layer modules in normal cortex and all CTE cases included in analyses.
