## Supplementary for "Spatially resolved transcriptomics reveals a unique disease signature and potential biomarkers for chronic traumatic encephalopathy": Figure S3.pdf

**Figure S3. Expression of CTE signature genes across brain cell types.** Shown are UMAP projections of expression profiles in aged brain, obtained from the Brain Knowledge Platform, for each of the 21 genes identified as upregulated in CTE lesions. The majority of the genes are expressed in astrocytes.

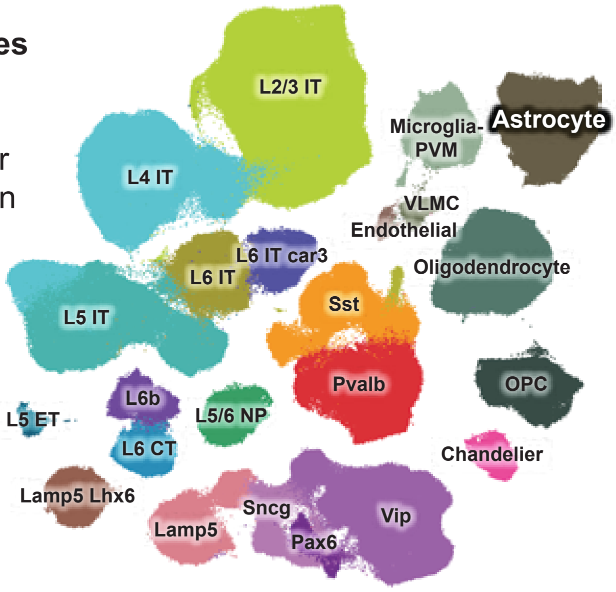


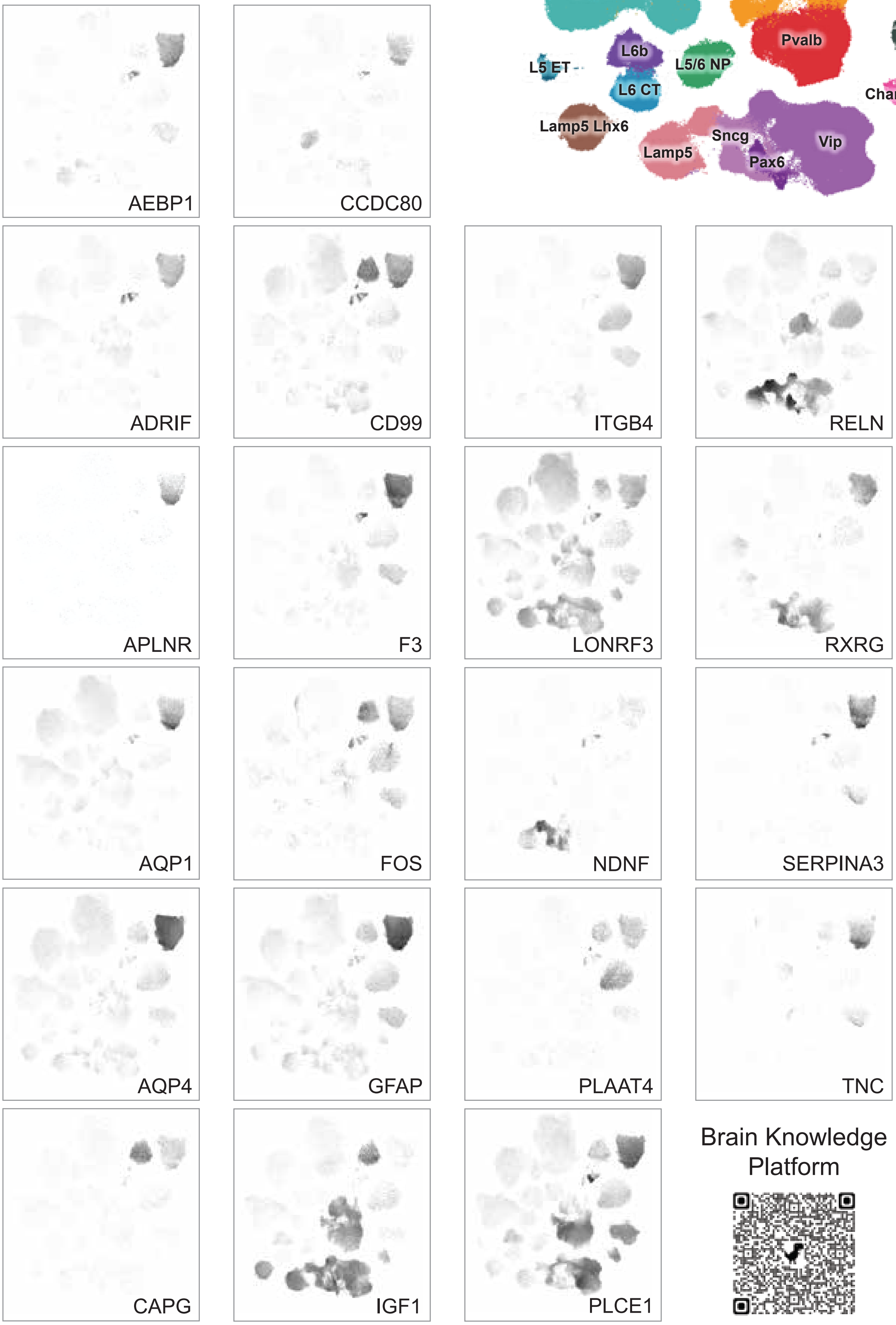
