## Supplementary for "Spatially resolved transcriptomics reveals a unique disease signature and potential biomarkers for chronic traumatic encephalopathy": Table S1.pdf

### Table S1: Cases used in this study and IHC summary

| Case | Decade at death | CTE stage* | Experiment** | AQP1 IHC | GFAP IHC |
|---|---|---|---|---|---|
| **1** | 4th | High (III-IV) | Visium, IHC | ↑ CTE DOS | ↑ CTE DOS |
| **2** | 6th | Low (II) | Visium, IHC | ↑ CTE DOS | ↑ CTE DOS, IAS |
| **3** | 5th | High (III) | Visium, IHC | ↑ CTE DOS, ↑ other DOS | ↑ CTE DOS, IAS |
| **4** | 6th | Low (I) | IHC | ↑ CTE DOS, ↑ other DOS | ↑ CTE DOS |
| **5** | 4th | Low (II-III) | IHC | ↑ CTE DOS | ↑ CTE DOS |
| **6** | 3rd | Low (I-II) | IHC | No abnormality | No abnormality |
| **7** | 5th | Low (I-II) | IHC | ↑ CTE DOS, ↑ other DOS | ↑ CTE DOS, IAS |
| **8** | 5th | High (III) | IHC | ↑ CTE DOS | ↑ CTE DOS, IAS |
| **control 1** | 5th | N/A | IHC | No abnormality | No abnormality |
| **control 2** | 3rd | N/A | IHC | No abnormality | No abnormality |
| **control 3** | 2nd | N/A | IHC | No abnormality | No abnormality |

* Current consensus stage (McKee stage); IHC, immunohistochemistry; DOS, depths of sulci; IAS, interface astroglial scarring

** A further individual was used for Visium transcriptional profiling, but did not pass sequencing QC so was excluded from all further analysis.
